# A human engineered mini-heart platform for mimicking ventricular pump function

**DOI:** 10.1101/2025.08.25.672056

**Authors:** Marcelo C. Ribeiro, Mariel Cano-Jorge, Simone ten Den, Danique Snippert, Marcel Karperien, Tom Kamperman, Guillaume Lajoinie, Michel Versluis, Robert Passier

## Abstract

Engineered cardiac tissue models for *in vitro* physiological studies often fail to replicate the pump function of the heart. Despite promising advancements, the use of engineered cardiac chambers is often hindered by complex fabrication processes and invasive characterization techniques. Here, we engineered a human cardiac chamber, referred to as a ‘mini-heart’, by employing a novel sacrificial molding approach within a customized bioreactor. The mini-heart’s pumping capability was confirmed through optical recording of fluid displacement at the engineered tissue inlet, enabling the non-invasive acquisition of hemodynamic parameters such as stroke volume, stroke work, ejection fraction, and developed pressure. Morphological analysis of the engineered tissues revealed organized sarcomeres and extracellular matrix self-determination, highlighting the advantage of our degradable mold technology. Additionally, we have measured calcium transients during both spontaneous and electrically-paced beating, and observed a positive inotropic response to the β-adrenergic agonist drug isoproterenol. Altogether, our biomimicking engineered tissue platform provides a robust tool for exploring cardiac pressure-volume dynamics, thereby facilitating complex disease modelling and drug screening applications *in vitro*.

## INTRODUCTION

Cardiovascular diseases (CVDs) remain the leading cause of death worldwide and are expected to increase in prevalence with global life expectancy (Tsao et al., 2023). Despite the urgent need to develop new drugs to treat CVDs, the output of the drug discovery pipeline has significantly declined in recent decades (Fordyce et al., 2015) primarily due to the lack of appropriate cardiac pre-clinical models(Kreutzer et al., 2022). Although animal models have served this purpose for decades, their efficacy remains questionable due to unreliable translation to human patients (Pound & Ritskes-Hoitinga, 2018). This highlights the need to develop more predictable and efficient human-based *in vitro* models that can faithfully recapitulate the healthy or diseased human heart, including its structural and functional complexity, as well as patient-specific phenotypes.

Since 2007, advancements in human pluripotent stem cell (hPSC) technology have facilitated the development of human-based models *in vitro* (Takahashi et al., 2007). Efficient and robust differentiation of hPSCs into cardiomyocytes (CMs) and other cardiac cell subpopulations has been established (Devalla & Passier, 2018) and used to create 3D cardiac tissues, such as engineered heart tissues (EHTs), which closely model the complexity of native heart tissue. EHTs demonstrate good predictive value for human (patho-)physiology (Sacchetto et al., 2020) and have been successfully explored for disease modelling and drug compound screening (Hansen et al., 2010; Nunes et al., 2013; Schwach et al., 2024). Despite their advantages, conventional EHTs also have limitations. These *in vitro* models typically consist of a strip of cardiac tissue formed around straight elastomeric supports or wires (Goldfracht et al., 2019; Nunes et al., 2013), consequently lacking the cavity that is necessary for fluid pumping function. This inability to direct fluid prevents them from replicating the heart’s pump function, thereby restricting their functional readouts to merely measuring twitch contractile forces. Moreover, conventional EHTs are often limited in their ability to independently and progressively fine-tune the cardiac preload and afterload imposed on the tissue due to dimensional constraints and the static properties of their supporting materials. This limitation results in an inaccurate mimicry and thus suboptimal study of the physiological dynamic stress experienced during ventricular systole and diastole as the progression of cardiovascular disease unfolds.

Recently, novel biotechnological methods have yielded engineered ventricular chambers capable of mimicking the heart’s pump function *in vitro*. As such, those models have the potential to facilitate the investigation of pressure and volume load dynamics in cardiac remodelling and disease *in vitro*. However, current fabrication approaches used to create these engineered ventricular chambers, including molding (R. A. Li et al., 2018), scaffold-seeding (Macqueen et al., 2018; Michas et al., 2022; Mohammadi et al., 2022; Patel & Birla, 2018) and bioprinting (Kupfer et al., 2020; Lee et al., n.d.) have significant limitations that hamper their use. One of the initial developments involved fabricating a scaffold-based ventricle using electrospun nanofibers, which was then seeded with cardiac cells (Macqueen et al., 2018). While this scaffold-based approach facilitated enhanced CM orientation, the rigidity and non-degradable nature of the scaffolds hampered cellular self-organization and matrix self-determination, preventing the cells from forming a cohesive tissue and impairing the full potential contractility of the cardiomyocytes. Alternatively, bioprinting of a cardiac chamber may offer a higher structural complexity. However, inherent limitations in the resolution of this technique often yield poor-quality dense cardiac constructs incapable of beating co-ordinately unless hPSCs are expanded and differentiated to CMs within a printed structure (Kupfer et al., 2020). Hence, development of engineered ventricles through molding techniques provides a balance between ease of fabrication, design complexity and cell remodelling accessibility. Nonetheless, a common limitation in the existing models is the need for multi-step manual handling of the preformed tissues, such as placing sutures (Macqueen et al., 2018), applying glue (Kupfer et al., 2020), or using folding mandrels (Mohammadi et al., 2022), to achieve successful fabrication and/or connection of perfusion tubes to the engineered chamber inlet. Apart from being laborious and complicated tasks, such handling introduces variability and may lead to improper sealing of the tissues, thereby hampering readouts and even failing studies. Furthermore, functional characterization of current engineered ventricles is often done utilizing conductance catheters (Kupfer et al., 2020; R. A. Li et al., 2018; Macqueen et al., 2018), which come with high costs, low accessibility and incompatibility with electrical pacing (Baan et al., 1985; Wei et al., 2014).

In this study, we developed a novel engineered pumping human heart chamber, referred to as a ‘mini-heart’. We employed an innovative approach using sacrificial molding that allowed for the straightforward fabrication, maintenance and characterization of the engineered tissue without requiring any direct manipulations of the cardiac tissue. Moreover, we evaluated the pump performance of the mini-heart via optical recording in a non-invasive manner, by measuring the volume displacement through a glass capillary connected to its lumen. This enabled the calculation of hemodynamic parameters such as stroke volume, stroke work, ejection fraction and developed pressures in the absence or presence of drug compounds and electrical stimulation. These results demonstrate the feasibility of reproducible manufacturing and non-invasive functional analysis of engineered ventricles, paving the way for the next generation of engineered pumping human heart chambers and opening new avenues for complex disease modelling, heart development and drug testing applications.

## RESULTS

### Fabrication of a mini-heart

To engineer a functional mini-heart capable of pumping fluid, we developed a novel sacrificial molding technique performed within a custom bioreactor. The bioreactor was designed to enable the fabrication, maintenance and acquisition of functional readouts of the mini-heart without requiring further manipulations. To achieve this, gelatin was cast around a ventricle-shaped PDMS mold inside the bioreactor. The mold was subsequently removed and replaced by a down-scaled replica made of gelatin and coupled to a glass capillary (Figure 1A). This established an ellipsoid-shell cavity between the two gelatin bodies, which was filled with hPSC-CMs and human cardiac fibroblasts (HCFs) embedded within a fibrin mix (Figure 1A iv,v). The mix polymerized into a fibrinogen matrix within minutes, casting the cells in the mini-heart shape imposed by the gelatin molds. The temperature was then raised to 37 °C enabling the thermal degradation and subsequent dissolution of the gelatin, leaving a single-inlet cardiac chamber internally and externally surrounded by culture medium (Figure 1C).

**Figure 1.**
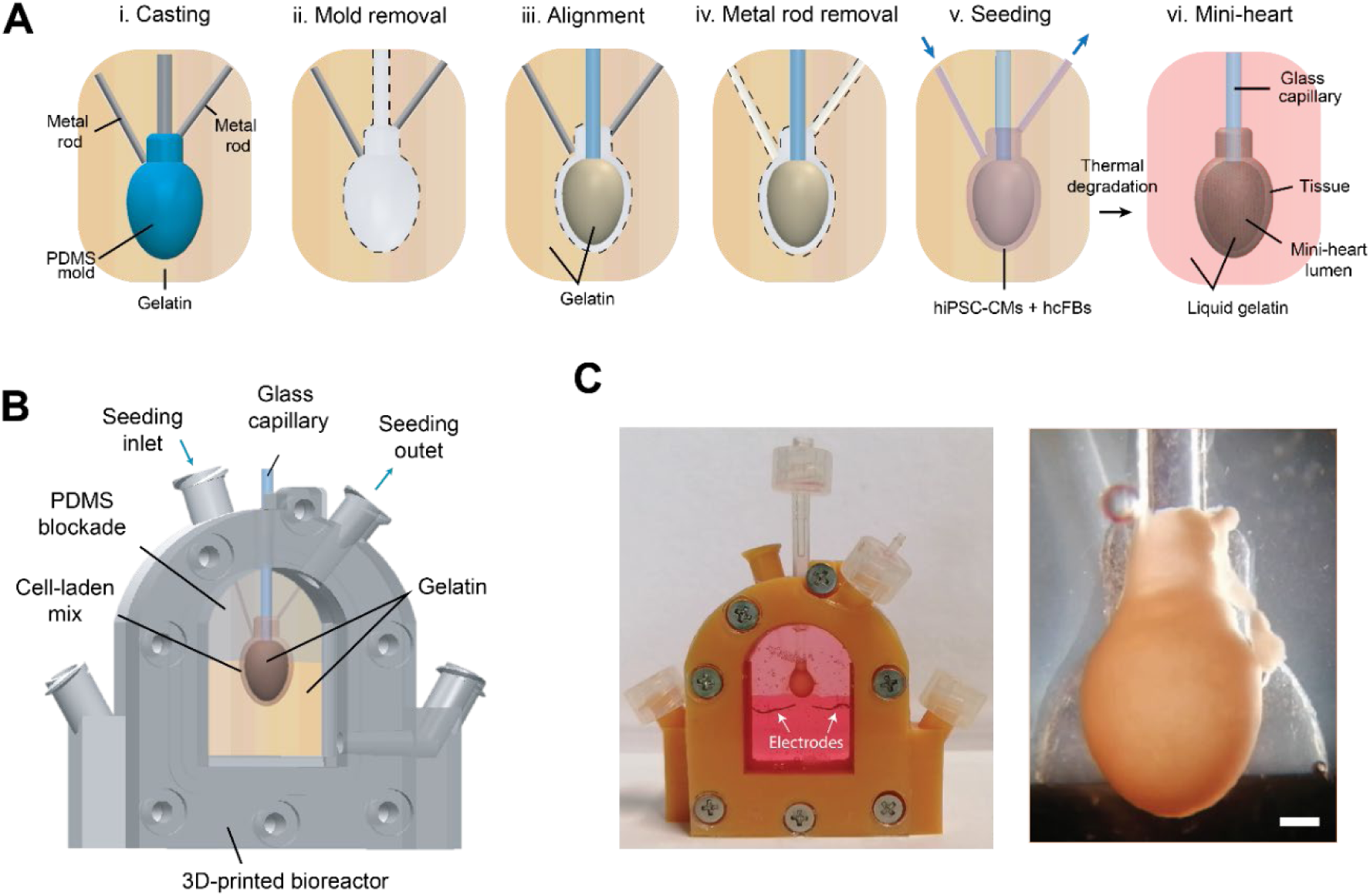
Fabrication of the mini-heart using a sacrificial moulding approach. A) i.) Casting of gelatin around PDMS mold and metal rods. ii) PDMS mold removal. iii) A smaller internal mold made of gelatin is added into the cavity left by the outer mold. iv) The metal rods are removed to create channels connecting the gap between the gelatin bodies. v) A cell-laden fibrinogen mix is seeded into the gap established by the gelatin molds. v) Thermal degradation of gelatin results into a hollow cardiac tissue. B) Diagram representing the seeding of the mini-heart inside a custom 3D-printed bioreactor. C) Bioreactor and mini-heart 10 days after formation. White arrows show pacing electrodes. *Scale bar* = 1mm.

Over time, cells migrate towards each other and form a compacted tissue (Figure 2C; Figure S4A), with estimated internal volumes of 88.16 ± 12.2 µL at day 1 (D1) to 31.15 ± 3.0 µL at day 20 (D20; Figure 2F). Importantly, this compaction induced the formation of impermeable tissue walls, as confirmed by the absence of fluorescent dextran outside the mini-heart when injected through the inlet of the chamber (Figure S4B). Analysis of cell viability of the mini-hearts revealed a significant decrease in cell viability from 92.83 ± 2.6 % at day 11 to 75.31 ± 6.1 % at day 20 (Figure 2A,B; Figure S3B). The wall thickness of the tissues at days 11, 15, and 20 was found to be 293.01 ± 48.7 µm, 283.57 ± 55.4 µm 276.21 ± 17.2 µm, respectively (Figure 2E; Figure S3A). Interestingly, immunohistochemistry analysis revealed the presence of highly organized sarcomeres (Figure 2D) restricted to 22.00 ± 1.41 µm of the outer core of the mini-hearts (at day 20). Towards the lumen of the tissues, less developed α-actinin positive cells were observed (Figure 2D). Positive expression of gap junction protein connexin-43 (Cx43) was found among hPSC-CMs, indicating electrical coupling (Figure 2I). Moreover, HCFs positive for vimentin were observed adjacent to hPSC-CMs and secreted collagen-1 matrix, providing a native cardiac ECM to the mini-heart (Figure 2H,J).

**Figure 2.**
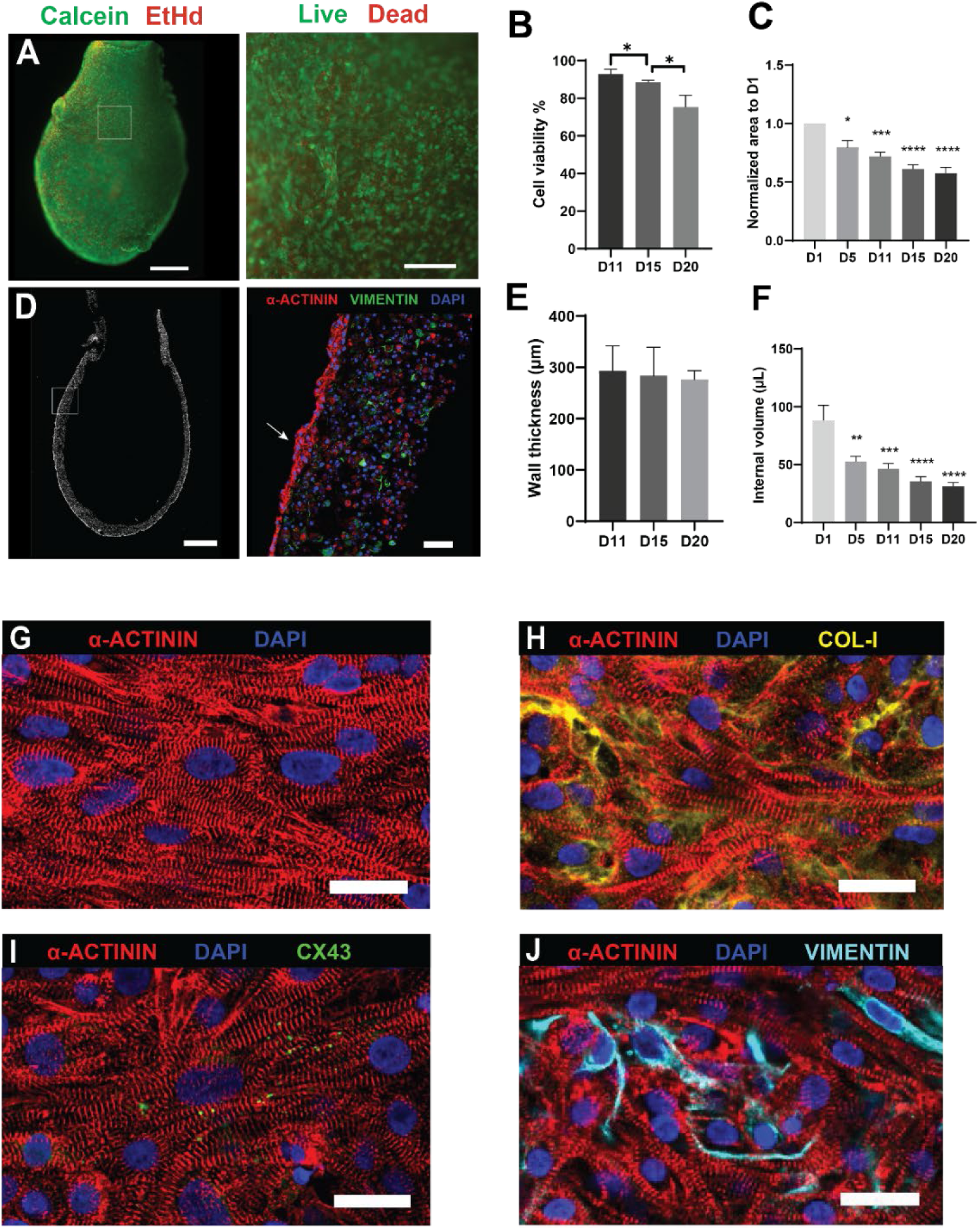
Morphological and histological analysis of mini-hearts. A) Viability live/dead assay. Scale bars: 1 mm (left) and 300 µm (right). Quantification of cell viability (B) and normalized area (C) of mini-hearts after fabrication. D) Cryosection of mini-heart and magnification of the wall depicting an organized sarcomere outer layer (arrow) (Scale bars= 1 mm (left); 50 µm (right)). Quantification of wall thickness (E) and estimation of mini-heart internal volume at days 1,5,11,15, and 20. G-J) Representative pictures of immunostaining for ɑ-actinin, collagen-I, connexin 43 and vimentin together with nuclear stain DAPI. Scale bars= 25 µm. Data plotted as means ± s.e.m., statistically tested by One-Way-ANOVA, (n=4 for wall thickness measurements; n=7, n=8 mini-hearts for normalized area and internal volume measurements, respectively). Significance was attributed to comparisons with values of P <0.05 *, P <0.01 **, P <0.001***, P<0.0001****.

### Non-invasive characterization of mini-heart pump performance

The main interest of developing engineered ventricles is to recapitulate the pump function of the heart *in vitro*. This results in a more physiological, organ-like model and provides clinically translatable readouts such as stroke volume, ejection fraction and developed pressure(Hsu et al., 2022; Konstam & Abboud, 2017). Therefore, we sought to evaluate the pump performance of our model in terms of these readouts. The mini-hearts exhibited spontaneous beating within 5 days post-fabrication (Supplementary Movie 1). From day 8, we observed an oscillating bidirectional flow in the glass capillary coupled to the lumen of the mini-heart (Figure 3A, Supplementary Movie 2). This flow was synchronized with the beating of the tissues, hereby confirming their pumping capability and enabling a non-invasive optical estimation of stroke volumes (Supplementary Movie 3, Figure 3B). To evaluate the pump performance over time, the mini-hearts were kept in culture for 20 days and stroke volumes were measured at days 11,15 and 20. Results indicated stroke volumes of 0.52 ± 0.1 µL, 0.99 ± 0.2 µL and 0.80 ± 0.2 µL, respectively (Figure 3C; Supplemental Movies 4-6). Based on the estimated internal volumes of the mini-hearts (Figure 2F), this corresponded to ejection fractions of 1.08 ± 0.1 %, 2.88 ± 0.5 %, and 2.57 ± 0.4 % for days 11,15 and 20, respectively (Figure 3D). The decrease in pump performance between days 15 and 20 may be linked to the decrease in tissue viability observed during this period.

**Figure 3.**
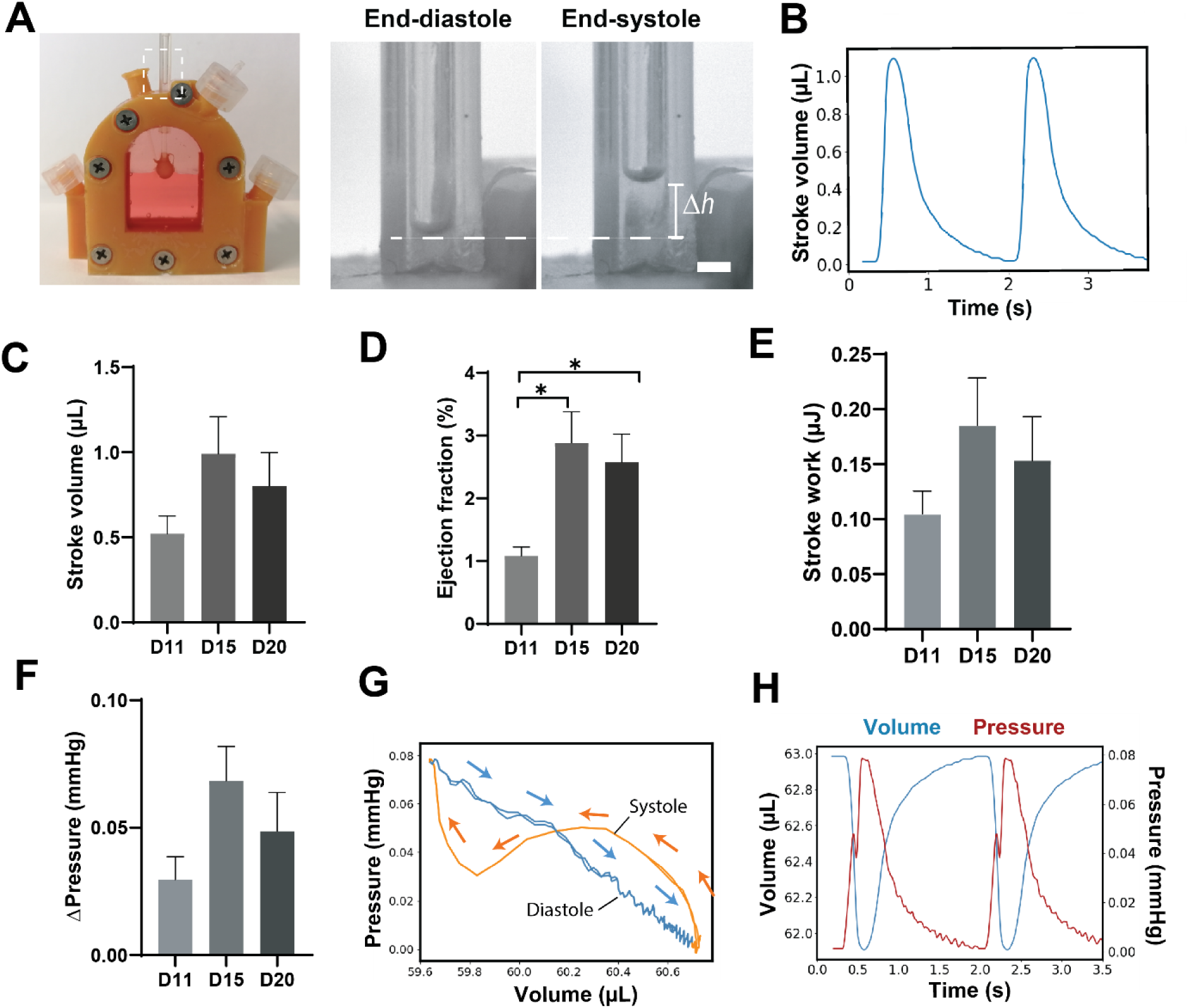
Non-invasive characterization of mini-heart pump performance. A) Displacement (*Δh*) of liquid within the glass capillary coupled to the lumen of mini-heart upon beating. B) Profile of stroke volume in time during spontaneous beating cycles. Stroke volume (C), ejection fraction (D), stroke work (E) and end-diastole to end-systole pressure difference (F) at days 11, 15 and 20. G) Pressure-volume diagram depicting the total volume inside the mini-heart and the corresponding pressure values inside the tissue. Systolic and diastolic phases indicated in orange and blue, respectively. H) Overlay of pressure and volume estimations in time during mini-heart beating cycle. Data plotted as means ± s.e.m., statistically tested by One-Way-ANOVA, (n=8 mini-hearts). * Denotes P <0.05.

Conductance catheters are typically used to measure internal pressure fluctuations within engineered ventricles (Wei et al., 2014). Although precise, this method is invasive, costly and incompatible with electrical pacing. We therefore estimated the pressure generated during the mini-heart’s beating cycles using an analytical model based on volume displacement at the tissue inlet. The total pressure (P_t_) developed by the mini-heart was defined as the sum of the hydrostatic pressure (P_h_), arising from fluctuations in the height of the liquid column, and the dynamic pressure (P_d_), due to the inertia of the liquid displaced during beating cycles (Figure 3G). This resulted in end-diastolic to end-systolic developed pressures ranging from 0.03± 0.01 mmHg to 0.07 ± 0.01 mmHg (Figure 3F). Although the total pressure is primarily governed by the hydrostatic component, as the contraction speed decreases approaching end-systole, the inertia of the fluid in the glass capillary increases the contribution of dynamic pressure to the overall pressure. This results in a slight drop in total pressure before end-systole is reached (Figure 3S). As anticipated, an inverse relationship between the total pressure P_t_ and the volume inside the mini-heart is observed (Figure 3H). However, the dynamic component P_d_ introduces a slight phase shift between the volume and pressure signals, resulting in a non-linear pressure-volume relationship (Figure 3G). The change in potential energy of the liquid displaced by the mini-heart in the glass capillary facilitated the estimation of the developed stroke works, with corresponding values of 0.10 ± 0.0 µJ, 0.18 ±0.1 µJ and 0.15 ± 0.0 µJ for days 11,15 and 20, respectively (Figure 3E).

To gain further insight into the performance of our heart model, we characterized the contractile kinetics of the mini-hearts based on the cross-section area dynamics during beating cycles (Figure 4A,B). The relative change between end-diastolic area (EDA) and end-systolic area (ESA) was used as an indicator of the extent of the contraction of the mini-hearts, which reached a peak value of 3.34 ± 0.5 % at day 15 (Figure 4C). A positive correlation trend (n.s.) was observed between the estimated ejection fractions and the relative EDA-to-ESA change (Figure S4C). The spontaneous beating frequency remained between 0.47 ± 0.1 Hz and 0.53 ± 0.1 Hz (Figure 4G). The cycle duration, relaxation time and relaxation velocities did not present any significant changes over time (Figure 4D,F,I). However, the time-to-peak decreased and contraction velocities (Figure 4E,H) increased significantly over time, suggesting an enhancement in hPSC-CM maturation.

**Figure 4.**
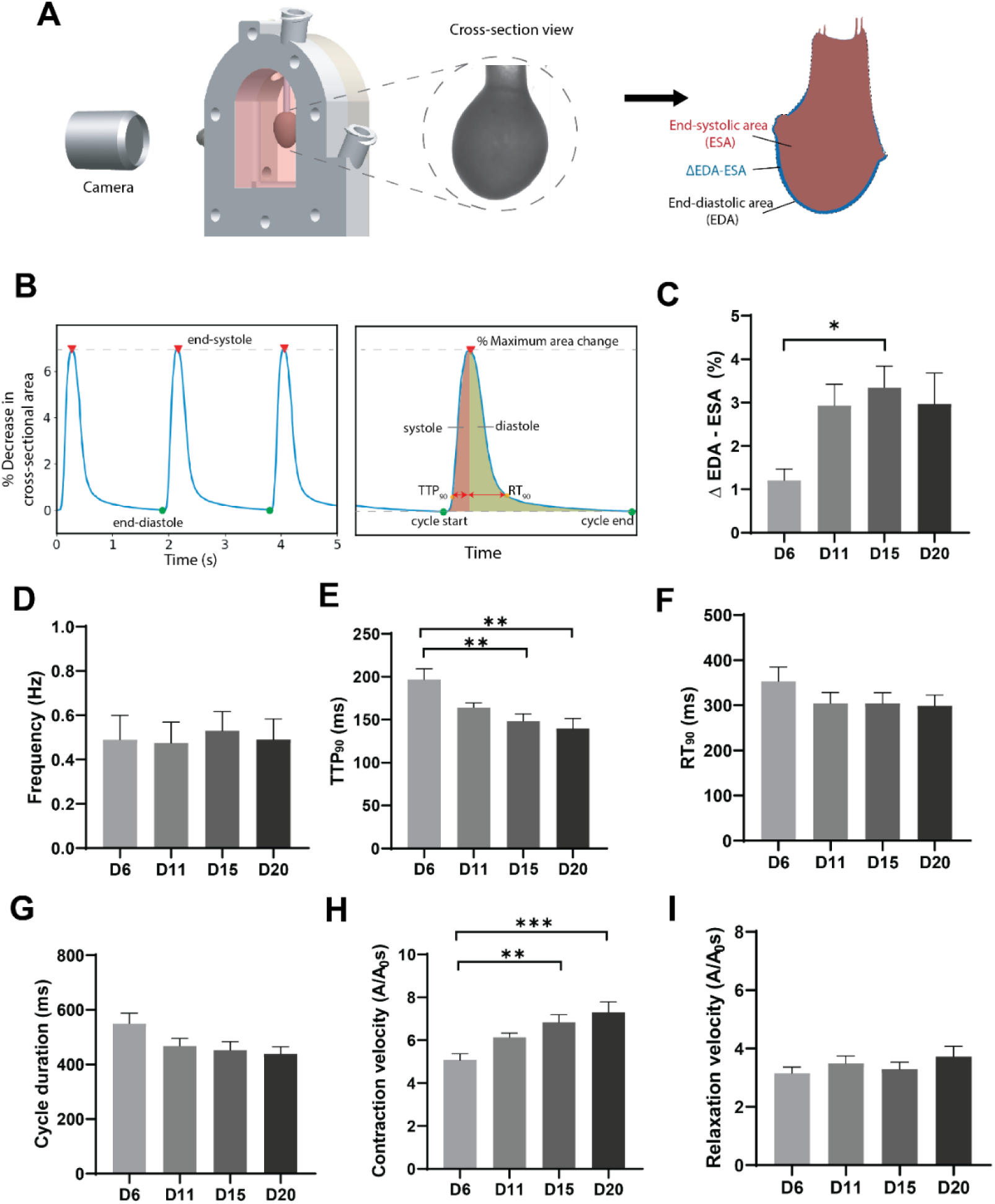
Analysis of contractile kinetics of mini-heart. A) Schematic depicting the imaging of end-systolic area (ESA) and end-diastolic area (EDA) of the mini heart. B) Diagram of the relative change in cross-section area in time with the corresponding associated contractile parameters. Evolution of relative EDA-to-ESA change (C), cycle duration (D), time to peak (TTP) (E), relaxation time (RT) (F), beating frequency (G), contraction and relaxation velocities (H,I) at days 6, 11, 15 and 20. Data plotted as means ± s.e.m., statistically tested by One-Way-ANOVA, (n=9 mini-hearts). Significance was attributed to comparisons with values of P <0.05 *, P <0.01 **, P <0.001***,

### Optical mapping for electrophysiological characterization of mini-heart

To assess the electrical activity of the mini-heart, we performed optical mapping of calcium transients. Isochronal maps enabled visualization of the calcium wave propagation, confirming the electrical coupling and stochastic source of depolarization of the mini-hearts (Figure 5A; Supplementary Movie 7). The tissues followed electrical pacing up to 3 Hz (Figure 5G). At pacing frequencies of 4 and 5 Hz, a decrease in contraction frequency was observed, likely due to impaired contractility resulting from insufficient recovery of CMs after fast depolarization cycles(Kanaporis & Blatter, 2015). However, this was accompanied by a significant decrease in the stroke volumes and the EDA-to-ESA relative change (Figure 5H,I), indicating a negative force-frequency relationship. This was further confirmed by a significant decrease in the peak amplitude of calcium transients for increasing pacing frequencies (Figure 5F). As anticipated, calcium transient duration (CTD_90_), relaxation times (RT) and time to peak (TTP) decreased with increasing pacing frequencies (Figure 5B-D). Although not significant, mean conduction velocities ranging from 1.58 ± 0.2 cm/s to 2.2 ± 0.6 cm/s presented an apparent increase proportional to the pacing frequency (Figure 5E).

**Figure 5.**
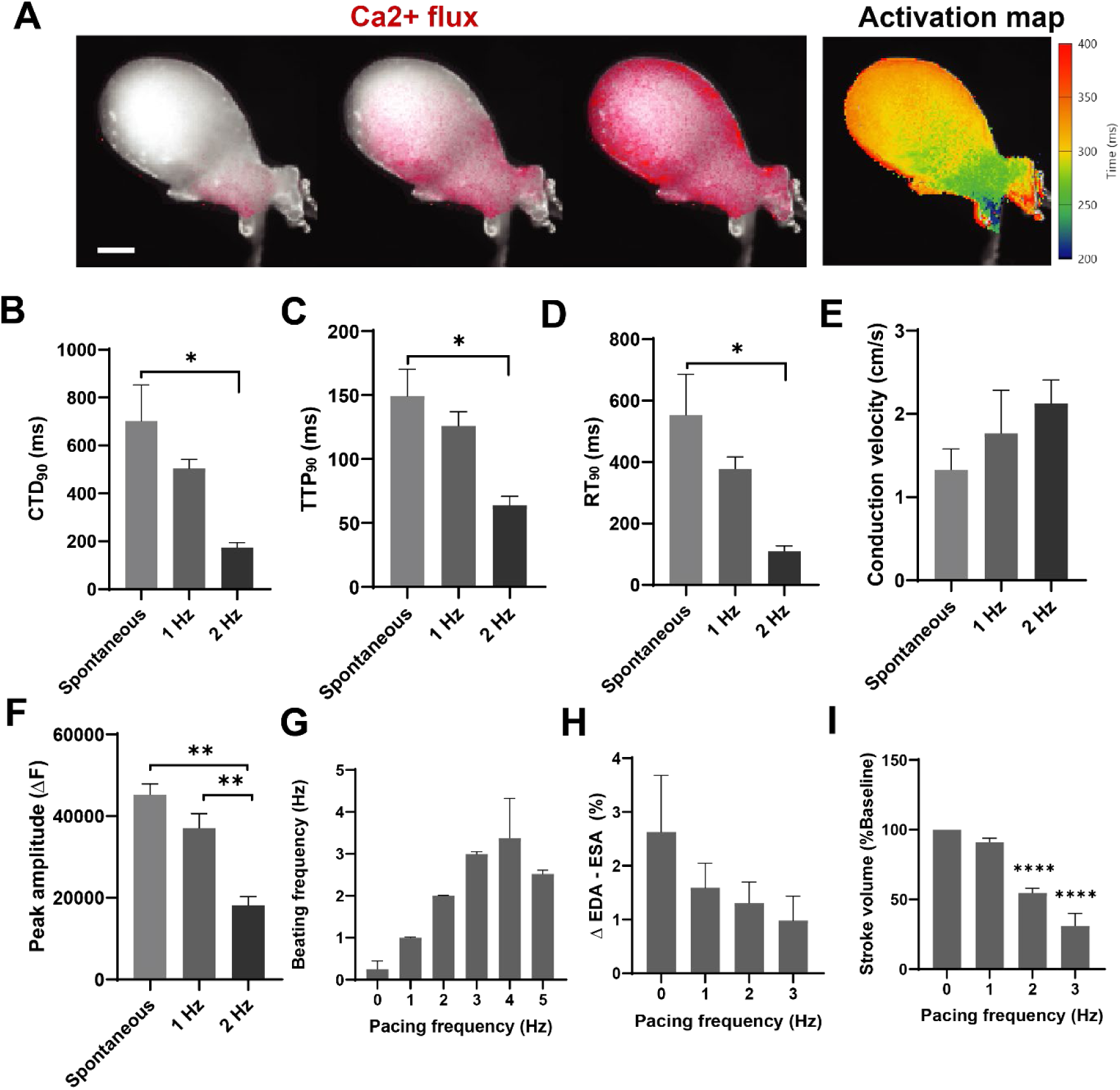
Optical mapping for electrophysiological characterization of mini-heart. A) Micrograph depicting the propagation of a calcium wave (red) though the mini-heart, with its corresponding activation map (Scale bar= 1mm). Quantification of calcium transient duration (CTD) (B), time-to-peak (TTP) (C), relaxation time (RT) (D), conduction velocity (E) and peak-amplitude (F) of mini-hearts beating spontaneously or electrically paced at 1 and 2 Hz. Electrical pacing of mini-heart at increasing frequencies (G) and the corresponding effects on (G), EDA-to-ESA change (H), and stroke volume (I). Data plotted as means ± s.e.m., statistically tested by One-Way-ANOVA, (n=4-5 mini-hearts). Significance was attributed to comparisons with values of P <0.05 *, P <0.01 **, P <0.001***, P<0.0001****.

### Mini-hearts exhibit functional responses to inotropic agents

Another hallmark of cardiac tissue functionality is their ability to respond to inotropic agents. Commonly, immature hPSC-CMs exhibit a positive chronotropic response to β-adrenergic stimulation but lack the positive inotropy characteristic of adult cardiomyocytes (Yang et al., 2014). We therefore evaluated the inotropic response of our mini-hearts to β-adrenergic agonist isoproterenol in concentrations ranging from 1 nM to 10 µM at a fixed pacing frequency of 1.5 Hz. A positive inotropic response was observed in both stroke volume and EDA-to-ESA measurements, which increased ∼50% and ∼180%, respectively, suggesting a significant level of maturation of hPSC-CMs in the mini-hearts (Figure 6A,C). The opposite effect was observed after administering the mini-hearts with increasing doses from 1 nM to 10 µM of calcium channel blocker nifedipine at a fixed pacing frequency of 1 Hz. As anticipated, stroke volumes and EDA-to-ESA readouts were attenuated with increasing concentrations of this compound (Figure 6B,C).

**Figure 6.**
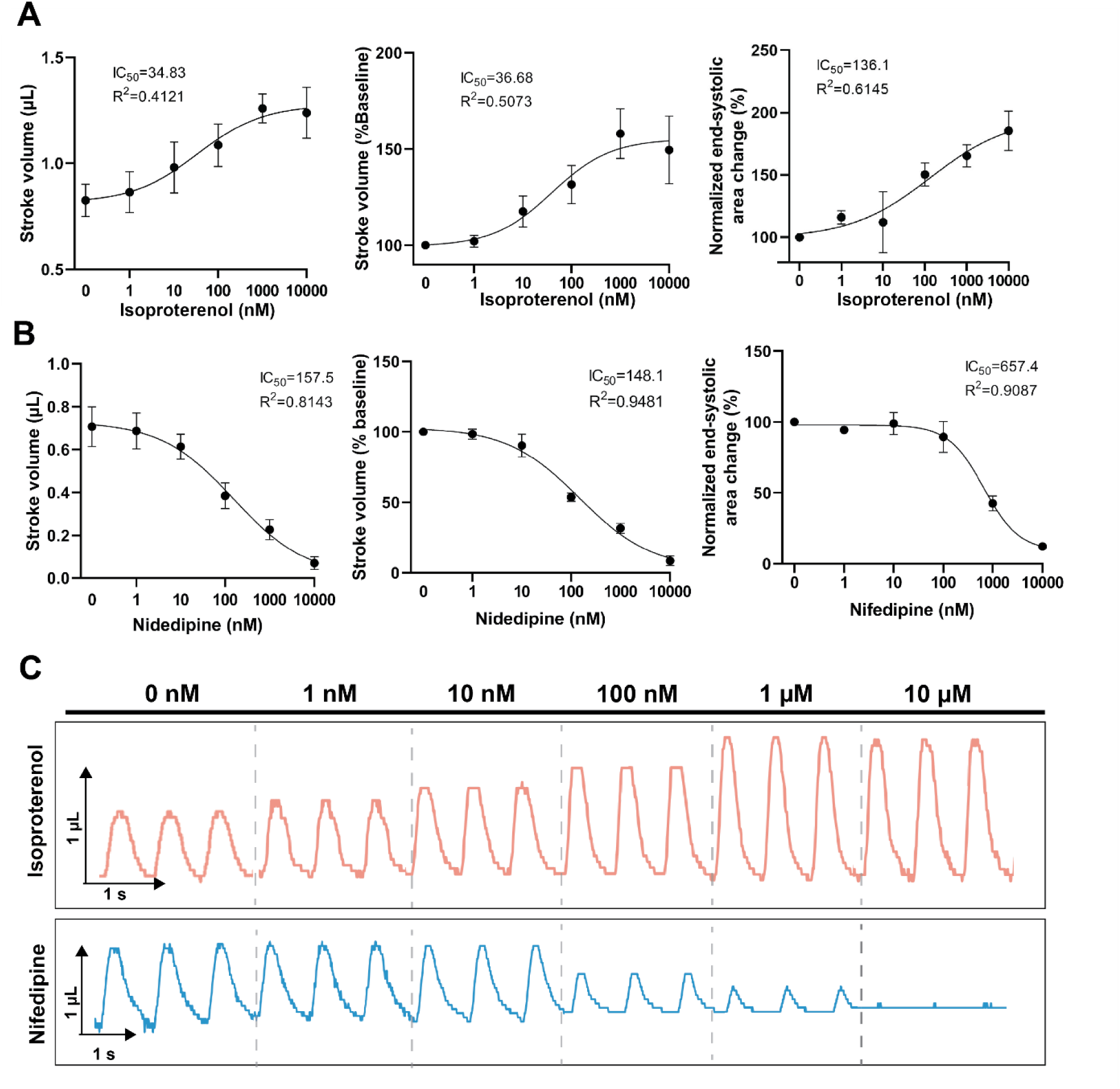
Response to drug compounds of mini-hearts. Stroke volumes and relative EDA-to-ESA change in response to cumulative concentration increase of isoproterenol (0-10 µM) (A), and nifedipine (0-10 µM). C) Schematic representation of stroke volumes in response to isoproterenol and nifedipine for two individual samples. Dashed lines separate responses to increasing drug concentrations within each sample, with a 15 minute incubation period per concentration. Values are expressed as means ± s.e.m. fitted to a sigmoidal curve, (n=4 mini-hearts for nifedipine, and n=5 for isoproterenol).

## DISCUSSION

In this study, we introduced a novel approach for efficiently creating hPSC-derived engineered ventricles that can generate cardiac pump function *in vitro.* Our novel fabrication approach utilizes thermally-degradable molds aligned within a custom bioreactor to facilitate the casting of a mini-heart. Consequently, our design enables proper attachment and sealing of the tissue around a glass capillary inlet, without the need for suturing or gluing, as described in previous studies (Kupfer et al., 2020; R. A. Li et al., 2018; Macqueen et al., 2018). Moreover, the fibrinogen matrix in which the cells are initially embedded undergoes complete degradation within a few days (Y. Li et al., 2015), which permits cellular self-organization and self-determination of ECM, a feature often limited in non-degradable scaffold-based constructs (Macqueen et al., 2018; Mohammadi et al., 2022). The dynamic self-organization of ECM components in our model likely facilitated the formation of compact, leak-free mini-hearts tightly enclosed around a glass capillary inlet, thereby achieving substantial ejection fractions. In turn, this allowed for visualizing liquid displacement in the glass capillary inlet during beating cycles, which made the non-invasive characterization of pump performance of the mini-hearts possible. This valuable feature highlights the accessibility and simplicity of our model, significantly increasing experimental success and reducing the costs and hurdles of conductance catheters that are commonly used to characterize pressure-volume dynamics in engineered ventricles. Nonetheless, a limitation of this approach is that pressure estimations are based on an analytical model where frictional components are neglected. Furthermore, due to the small size of the glass capillary, it is possible that part of the pumping power is dissipated within the heart itself, which could result in the underestimation of recorded pressure measurements at the glass capillary inlet compared to the actual pressure generated. Regardless of the assumptions on the absolute values, these pressure measurements may be used for relative comparisons between experimental groups in disease modelling and drug testing, thus inferring clinical significance to *in vitro* settings.

The presence of valves in human ventricles facilitates isovolumetric contraction and relaxation that result in characteristic clinical pressure-volume (PV) loop patterns. In contrast, valve-less engineered cardiac chambers exhibit less distinct delays between pressure and volume fluctuations during beating cycles, typically leading to narrower, elliptical PV-loop patterns (R. A. Li et al., 2018; Macqueen et al., 2018). In our model, we observed a non-linear PV-relationship characterized by a double-helix pattern. This unique behaviour can be attributed to the incorporation of a fluidic column at the tissue inlet, which influences the pressure dynamics within the tissue, unlike most other valve-less cardiac chambers that operate in isolation. Nonetheless, this distinct PV-relationship patterns offer valuable insights into the asymmetries between contractile and relaxation profiles. Since relaxation velocities were generally lower than contraction velocities in our model, inertial effects during systole contributed more significantly to the total pressure than during diastole. This disparity is evident in the pressure-volume relationship, where the systolic branch exhibits a non-linear behaviour due to stronger fluctuations in dynamic pressure, while the diastolic branch of the curve displays an almost linear pattern dominated by the hydrostatic pressure. Although the clinical interpretation is not straightforward, this visualization can help identify overall changes in contractile or relaxation velocities, with increased velocities reflected as wider PV-loop relationships, and decreased velocities resulting in more linear-like patterns.

In terms of performance, our mini-hearts displayed spontaneous beating with frequencies, APD, TTP and RT values comparable to other engineered ventricles (R. A. Li et al., 2018). Although far from human ejection fraction values (50%-70%), the ejection fractions of our mini-heart (∼1%-3%) are in the same range as other reported models (0.2%-9.6%) (Kupfer et al., 2020; R. A. Li et al., 2018; Macqueen et al., 2018; Michas et al., 2022). The mini-hearts demonstrated stroke works of ∼0.10 - ∼0.18 µJ which, to our knowledge, are the highest stroke work values compared to other engineered ventricles (6 nJ-57 nJ) (Kupfer et al., 2020; R. A. Li et al., 2018; Macqueen et al., 2018). Moreover, we observed a positive inotropic response to isoproterenol, reaching a ∼50% increase in stroke volume for a 1 µM dose, and a EC_50_ of ∼36 nM. This is the highest force gain reported so far, and the lowest EC_50_, with respect to a similar model featuring a ∼10% increase in stroke volume and a EC_50_ of 237 nM (for cardiac output) in response to isoproterenol (R. A. Li et al., 2018). These relatively high-performance outputs reiterate the significance of our innovative fabrication method in advancing the modelling of the human heart *in vitro*. Together with the decrease in TTP observed over culture time, these findings suggest enhanced cardiomyocyte maturation of the mini-hearts. However, the observed negative force-frequency relationship is consistent with findings in similar models (R. A. Li et al., 2018; Macqueen et al., 2018) and suggests the calcium handling of hPSC-CMs still requires further maturation.

Increase in cardiac wall thickness is commonly associated with enhanced pump performance. Although our mini-hearts had a wall thickness of ∼275 µm at day 20, we observed an outer core of elongated CMs with organized sarcomeres restricted to ∼22 µm. Since this CM layer was responsible for leading the synchronous contraction of the mini-heart, a follow-up study should explore the incorporation of enhanced perfusion strategies, and tissue vascularization, to increase the thickness of this organized CM layer, thereby achieving more physiological ejection fraction values. Although human-engineered cardiac chambers hold potential to develop more accurate *in vitro* models, the low throughput associated with their fabrication and characterization remains a challenge. In this regard, the time and cost effectiveness of our functional model, will promote the accessibility and standardization of engineered heart chambers. We believe this will facilitate the overarching goal of establishing accurate, patient-specific *in vitro* disease models, ultimately enabling more predictable modelling of human heart diseases and improving drug efficacy screening early in drug discovery. In combination with efforts in enhancing the throughput and maturation of hPSC cardiac lineages, these advances will also bring us a step closer to regenerative medicine applications.

## Online Methods

### HPSC Culture and Generation of hPSC-CMs

The experiments were performed using human-induced pluripotent stem cells (hPSCs; Coriell, GM25256) derived cardiomyocytes (CMs) and human cardiac fibroblasts (HCFs) (Promocell, C-12375). Differentiation to hPSC-CMs was induced as described previously (Birket et al., 2015). Briefly mesodermal differentiation was induced by Activin-A (20-30 ng/mL, Miltenyi, Leiden, The Netherlands, 130–115-010), BMP4 (20 ng/mL, R&D systems, Minneapolis, MN, USA, 314-BP/CF), and WNT activator CHIR99021 (1.5-2.25 μmol/L, Axon Medchem, Groningen, The Netherlands, 1386) in BPEL medium(Ng et al., 2008). At day 3, cells were refreshed with BPEL containing WNT inhibitor XAV939 (5 μmol/L, R&D Systems 3748). The cells were maintained in our previously described cardiomyocyte medium (CM medium; (Birket et al., 2015) with additional 5 mM sodium DL-lactate solution (60%, Sigma Aldrich, cat. no. L4263), to achieve metabolic purification of hPSC-CMs. The cells were then dissociated with TrypLE 10X (ThermoFisher, A1217702) and analyzed by FACS to ensure >80% positive expression of cardiac troponin T (cTnT) prior to cryopreservation.

### Culture of HCFs

HCFs were expanded according to the manufacturer’s instructions. Briefly, HCFs were thawed and maintained in FGM-3 medium (PromoCell, C-23025). Refreshments were done every 48 hours until 70-90% confluency was reached. The cells were passaged and expanded until reaching 11 passages, prior to cryopreservation.

### Fabrication of the platform and mini-heart formation

The tissues were generated using a sacrificial molding approach (Figure 1). A custom-made bioreactor was designed in Solidworks (2019) and 3D-printed (Rapidshape S30L) with a biocompatible resin (MP300 500GM, Harshad). The bioreactor is composed of two parts that can be assembled using bolts and nuts. Each part hosts an imaging window made from PMMA (Merck GF53167608; 1 mm thick) and two luer-lock-compatible inlets (Figure S1A). One of the parts features a PDMS blockade designed to guide the alignment of inserts. Next, a custom-shaped PDMS (10:1; Mavom 1060040) mold was placed inside the bioreactor (Figure S1C), which was subsequently closed. Two metal rods (1 mm diameter) were then added, connecting the mold with the two upper luer-lock inlets (Figure 1A). Gelatin (Sigma-Aldrich G1890; 10%) was prepared in our CM medium and was casted around the mold in two steps. First, the bioreactor was placed horizontally and half-filled with liquified gelatin. After solidification of gelatin at 4 °C, the remaining half was added and cooled until solidified (Figure S1D). This sequential gelation creates two independent gelatin bodies within the bioreactor, allowing it to be opened without damaging the casted gelatin. The PDMS mold was then removed from the bioreactor (FigureS1E) and replaced with a down-scaled replica made of gelatin (inner-mold) (Figure S1F), which was coupled to a glass capillary (World Precision Instruments, TW150-3) and a supporting tubing to prevent it from sliding (Figure S2). The inner-mold was made by filling a negative PDMS mold with liquified gelatin through the glass capillary inlet and subsequently freezing it. Finally, the bioreactor was re-closed. Since both gelatin parts were originally casted within the closed bioreactor, they create a seamless seal upon reassembly, ensuring mold accuracy. The metal rods were pulled out through the luer lock inlets to create two channels connected to the ellipsoid shell cavity formed between the gelatin bodies (Figure S2G,H).

Next, hPSC-CMs were thawed, counted and resuspended in CM medium together with HCFs at 10:1 ratio. An extracellular matrix (ECM) solution consisting of 2X CM medium, fibrinogen (final concentration of 2 mg/mL, Sigma-Aldrich F8630), Matrigel (final concentration of 1 mg/mL), and aprotinin (final concentration of 2.5 µg/mL, Sigma-Aldrich, A1153) was prepared on ice and added to the cell suspension to yield a final concentration of 20 x 10^6^cells/mL. Next, 0.6 U/mL of thrombin (Sigma, T7513) was added to the solution. The mix was quickly loaded into the seeding channel of the bioreactor to fill the cavity between the gelatin bodies (Figure 1B). The seeded bioreactor was incubated at RT for 10 mins to ensure fibrinogen polymerization. Next, the bioreactor was incubated at 37°C for 30 min to induce thermal degradation of gelatin. After this time, the supporting tubing was removed from the lumen of the chamber. The formed tissue was maintained at 37°C and 5% CO_2_. After 24 h, the liquified gelatin was replaced with CM medium. Refreshments were done daily and after electrical pacing. The PDMS blockade remained in the bioreactor without interfering with the formed tissue. One of the luer-lock inlets of the bioreactor remained opened during tissue maintenance, allowing for continuous medium gas exchange.

### Functional analysis

The non-invasive acquisition of readouts was based on the optical tracking of the mini-heart cross sectional area and the bidirectional displacement of liquid through the glass capillary coupled to its inlet. To achieve this, a digital camera (a2A4200-40umPRO, Basler ace; lens LM50FC24M, Kowa) was positioned in front of the bioreactor to record the tissues at 100 fps with a custom acquisition program made in LabVIEW (2019). The videos were processed using custom ImageJ macro and Python scripts. Stroke volumes were estimated as the cylinder volume corresponding to the length of liquid displaced through the glass capillary with internal diameter of 1.12 mm. The averaged displacement of liquid over 3 beating cycles was used to estimate the stroke volume at each measurement timepoint. The volume of the mini-heart was estimated from cross-section area measurements as an ellipsoid cap assuming an equal semi-minor axis perpendicular to the imaging plane. The stroke work (*SW*) was calculated as the change in potential energy of the volume of liquid in the glass capillary (V_c_) vertically displaced by a distance Δℎ (from end-systole to end-diastole; Figure 3A) as *SW* = *V*_*C*_ *ρg*Δℎ, with ρ=993.37 kg/m^3^ approximated as the density of liquified gelatin, and *g* the gravitational constant.

The pressure generated during mini-heart contractions was estimated based on the temporal position (*h(t)*) of the liquid level in the glass capillary coupled to the tissue inlet. The total pressure developed (*P_t_*) within the tissue was analytically estimated accounting for hydrostatic (*P_h_*) and dynamic pressure (*P_d_*) components, such that:

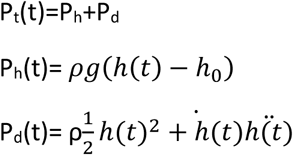

where *h_0_*, is the level of liquid at end-diastole, *ρ*=993.37 kg/m^3^ approximated as the density of liquified gelatin, and *g* the gravitational constant. The developed pressure per beat *ΔP* was taken as the difference between end-diastolic and end-diastolic pressures and was acquired at days 11,15 and 20.

The baseline volume of the tissue at end-diastole was estimated from its cross-section area as previously described. The volume as a function of time was obtained by subtracting the stroke volume in time from the baseline volume.

Responses to drug compounds were evaluated at day 15. For a positive inotropic response, the tissues were exposed to increasing cumulative concentrations (1 nM, 10 nM, 100 nM, 1µM, 10 µM) of β - adrenergic agonist isoproterenol (Sigma, I5627). Negative inotropic effects were assessed with administration of increasing doses (1 nM, 10 nM, 100 nM, 1µM, 10 µM) of the calcium channel blocker nifedipine (Sigma, N7634). The response to both drugs were calculated relative to the base-line measurement. Stroke volumes and cross sectional areas were measured 15 minutes after each dose administration to ensure uniform dilution of the drug inside the bioreactor.

### Calcium imaging and electrical pacing

For optical mapping of calcium propagation, the tissues were incubated for 1 hour at 37 °C in a phosphate-buffered saline (PBS) solution containing 5 µM calcium dye Fluo-8 (Abcam, AB142773), 1% (vol/vol) pluronic (Sigma-Aldrich P2443) in Bovine Serum Albumin (New England Biolabs B9000S) and 15 mM glucose at day 14. After one wash in PBS, the samples were mounted in a glass-bottom dish (Ibidi 81158) in CM medium and imaged under a Nikon Ti2-E inverted microscope at 50 fps. The recordings were processed with ImageJ and analyzed with BV Workbench 2.7.2 The tissues were electrically paced at 1 Hz for 30 s (10 ms biphasic pulses, 2 V/cm) with a custom-made voltage-source connected to 2 platinum electrodes placed approximately 6 mm away from the apex of the tissue and 10 mm apart from each other. In frequency pacing experiments, the tissues were paced at 1,2,3,4, and 5 Hz using the previously described settings.

### Immunostaining and histology

The mini-hearts were fixed in 4% paraformaldehyde in PBS for 45 min at room temperature. For whole-mount staining, tissues were washed three times in PBS, permeated with 0.3% Triton-X 100 (Sigma-Aldrich) for 20 min and blocked in 3% BSA, 0.3% Triton-X 100, and 0.1% Tween in PBS overnight at 4 °C. Next, the tissues were incubated for 2 days at 4 °C with primary antibody anti-alpha-actinin (1:400; Invitrogen, MA5-12960), anti-vimentin (1:200; Novus Biologicals NBP1-85814), anti-connexin-43 (1:200; Sigma-Aldrich, C6219) or anti-collagen-I (1:200; Novus Biologicals, NB600-408). After three washes in 0.3% Triton-X 100 (3 × 20 min), the samples were incubated with secondary antibody Goat-anti-Mouse IgG Alexa Fluor 647 (1:500; Invitrogen, A21235), and/or Goat-anti Rabbit IgG Alexa Fluor 488 (1:500; Invitrogen, A27034) and DAPI for 24 h at 4 °C. Samples were washed three times in PBS, mounted on a glass-bottom plate (Ibidi 81158) and imaged with a Zeiss LSM 880 confocal microscope.

For cryosections, the samples were fixed as previously mentioned and incubated overnight in a 30% sucrose solution in PBS. The tissues were embedded in CryoMatrix Gel (Thermo Scientific, 6769006) and cryopreserved by immersion in 2-methylbuthane. Next, 12 µm sections were obtained in an MNT (SLEE Medical) cryostat. The sections were permeated with 0.1% Triton-X 100 (Sigma-Aldrich) in PBS for 8 min and blocked with 1% (*vol/vol*) BSA in PBS for 1 h. Next, the sections were incubated overnight at 4 °C with primary antibody anti-alpha-actinin (1:800; Sigma, A7811) and anti-vimentin (1:200; Novus Biologicals NBP1-85814). Sections were washed three times in PBS and incubated with secondary antibody Goat-anti-Mouse IgG Alexa Fluor 647 (1:500; Invitrogen, A21235) and Goat-anti Rabbit IgG Alexa Fluor 488 (1:500; Invitrogen, A27034) for 1 h at room temperature. Sections were washed three times in PBS and stained with DAPI. The samples were mounted with ProLong Gold antifade (Life Technologies) and imaged with a Zeiss LSM 880 laser confocal microscope.

For live/dead staining, tissues were incubated in PBS with 3 µM Calcein and 4 µM Etdh1 (LIVE/DEAD; Invitrogen L3224) for 45 min at 37 °C. The samples were washed in PBS and imaged under a Nikon Ti2-E inverted microscope. All images were analyzed with ImageJ.

Tissue permeability was assessed by injecting fluorescently-labelled dextran (40 kDa, Fisher Scientific, D1845) into the lumen of the mini-heart via the glass capillary inlet. The mean fluorescence intensity of the tissue and the surrounding background were quantified over time using ImageJ.

### Statistics

Statistical analysis was performed using GraphPad Prism 8. Results are reported as the mean ± standard error of the mean (s.e.m). One-way ANOVA tests were done to compare differences between timepoints or pacing frequency measurements. Significance was attributed to comparisons with values of P <0.05 *, P <0.01 **, P <0.001***, P<0.0001****.

## Acknowledgements

The authors acknowledge Robert Molenaar for his technical support regarding the imaging setup, and Charlotte Nawijn and Macy Vreman for their valuable support with the printing of bioreactors. R.P. acknowledges financial support from the European Research Council (ERC, Advanced Grant Heart2Beat, project number 101098372). M.C.R. acknowledges 101070953 — BioRobot-MiniHeart — HORIZON-EIC-2021-PATHFINDERCHALLENGES-01. M.C.-J. and R.P. acknowledge NWO through grant NWO/ENW-XL 2019.029.

## Author contributions

M.C.R., M.C.-J., G.L., and R.P. conceived and designed the idea. M.C.-J. collected the data. M.C.R., M.C.-J., and R.P. analyzed and interpreted the data. M.C.-J., M.C.R., and R.P. wrote the manuscript. All authors have reviewed and approved the manuscript.

## Data availability statement

The data that support the findings of this study are available from the corresponding author upon reasonable request.

## Glossary

CVD: Cardiovascular disease
hPSC: Human pluripotent stem cell
CM: Cardiomyocyte
EHT: Engineered heart tissue
HCF: Human cardiac fibroblast
Cx-43: Connexin 43
ECM: Extracellular matrix
P_t_: Total pressure
P_d_: Dynamic pressure
P_h_: Hydrostatic pressure
EDA: End-diastolic area
ESA: End-systolic area
CTD: Calcium transient duration
RT: Relaxation time
TTP: Time to peak
PV: Pressure-volume
cTnT: Cardiac troponin
SW: Stroke work

## References

Baan, J., Van Der Velde, T., De Bruin, H. G., Smeenk, G. J., Koops, J., Van Dijk, A. D., Temmerman, D., Senden, J., & Buis, B. (1985). Continuous measurement of left ventricular volume in animals and humans by conductance catheter. Circulation, 70(5), 812–823. http://ahajournals.org

Birket, M. J., Ribeiro, M. C., Kosmidis, G., Ward, D., Leitoguinho, A. R., van de Pol, V., Dambrot, C., Devalla, H. D., Davis, R. P., Mastroberardino, P. G., Atsma, D. E., Passier, R., & Mummery, C. L. (2015). Contractile Defect Caused by Mutation in MYBPC3 Revealed under Conditions Optimized for Human PSC-Cardiomyocyte Function. Cell Reports, 13(4), 733–745. 10.1016/j.celrep.2015.09.025

Devalla, H. D., & Passier, R. (2018). Cardiac differentiation of pluripotent stem cells and implications for modeling the heart in health and disease. In Sci. Transl. Med (Vol. 10). http://stm.sciencemag.org/

Fordyce, C. B., Roe, M. T., Ahmad, T., Libby, P., Borer, J. S., Hiatt, W. R., Bristow, M. R., Packer, M., Wasserman, S. M., Braunstein, N., Pitt, B., Demets, D. L., Cooper-Arnold, K., Armstrong, P. W., Berkowitz, S. D., Scott, R., Prats, J., Galis, Z. S., Stockbridge, N., … Califf, R. M. (2015). THE PRESENT AND FUTURE STATE-OF-THE-ART REVIEW Cardiovascular Drug Development Is it Dead or Just Hibernating*?*

Goldfracht, I., Efraim, Y., Shinnawi, R., Kovalev, E., Huber, I., Gepstein, A., Arbel, G., Shaheen, N., Tiburcy, M., Zimmermann, W. H., Machluf, M., & Gepstein, L. (2019). Engineered heart tissue models from hiPSC-derived cardiomyocytes and cardiac ECM for disease modeling and drug testing applications. Acta Biomaterialia, 92, 145–159. 10.1016/j.actbio.2019.05.016

Hansen, A., Eder, A., Bönstrup, M., Flato, M., Mewe, M., Schaaf, S., Aksehirlioglu, B., Schwörer, A., Uebeler, J., & Eschenhagen, T. (2010). Development of a drug screening platform based on engineered heart tissue. Circulation Research, 107(1), 35–44. 10.1161/CIRCRESAHA.109.211458

Hsu, S., Fang, J. C., & Borlaug, B. A. (2022). Hemodynamics for the Heart Failure Clinician: A State-of-the-Art Review. In Journal of Cardiac Failure (Vol. 28, Issue 1, pp. 133–148). Elsevier B.V. 10.1016/j.cardfail.2021.07.012

Kanaporis, G., & Blatter, L. A. (2015). The mechanisms of calcium cycling and action potential dynamics in cardiac alternans. Circulation Research, 116(5), 846–856. 10.1161/CIRCRESAHA.116.305404

Konstam, M. A., & Abboud, F. M. (2017). Ejection Fraction: Misunderstood and Overrated (Changing the Paradigm in Categorizing Heart Failure). Circulation, 135(8), 717–719. 10.1161/CIRCULATIONAHA.116.025795

Kreutzer, F. P., Meinecke, A., Schmidt, K., Fiedler, J., & Thum, T. (2022). Alternative strategies in cardiac preclinical research and new clinical trial formats. In Cardiovascular Research (Vol. 118, Issue 3, pp. 746–762). Oxford University Press. 10.1093/cvr/cvab075

Kupfer, M. E., Lin, W. H., Ravikumar, V., Qiu, K., Wang, L., Gao, L., Bhuiyan, D. B., Lenz, M., Ai, J., Mahutga, R. R., Townsend, D., Zhang, J., McAlpine, M. C., Tolkacheva, E. G., & Ogle, B. M. (2020). In Situ Expansion, Differentiation, and Electromechanical Coupling of Human Cardiac Muscle in a 3D Bioprinted, Chambered Organoid. Circulation Research, 127(2), 207–224. 10.1161/CIRCRESAHA.119.316155

Lee, A., Hudson, A. R., Shiwarski, D. J., Tashman, J. W., Hinton, T. J., Yerneni, S., Bliley, J. M., Campbell, P. G., & Feinberg, A. W. (n.d.). 3D bioprinting of collagen to rebuild components of the human heart. http://science.sciencemag.org/

Li, R. A., Keung, W., Cashman, T. J., Backeris, P. C., Johnson, B. V., Bardot, E. S., Wong, A. O. T., Chan, P. K. W., Chan, C. W. Y., & Costa, K. D. (2018). Bioengineering an electro-mechanically functional miniature ventricular heart chamber from human pluripotent stem cells. Biomaterials, 163, 116–127. 10.1016/j.biomaterials.2018.02.024

Li, Y., Meng, H., Liu, Y., & Lee, B. P. (2015). Fibrin gel as an injectable biodegradable scaffold and cell carrier for tissue engineering. In Scientific World Journal (Vol. 2015). Hindawi Limited. 10.1155/2015/685690

Macqueen, L. A., Sheehy, S. P., Chantre, C. O., Zimmerman, J. F., Pasqualini, F. S., Liu, X., Goss, J. A., Campbell, P. H., Gonzalez, G. M., Park, S. J., Capulli, A. K., Ferrier, J. P., Fettah Kosar, T., Mahadevan, L., Pu, W. T., & Parker, K. K. (2018). A tissue-engineered scale model of the heart ventricle. Nature Biomedical Engineering, 2(12), 930–941. 10.1038/s41551-018-0271-5

Michas, C., Çağatay Karakan, M., Nautiyal, P., Seidman, J. G., Seidman, C. E., Agarwal, A., Ekinci, K., Eyckmans, J., White, A. E., & Chen, C. S. (2022). Engineering a living cardiac pump on a chip using high-precision fabrication. In Sci. Adv (Vol. 8).

Mohammadi, M. H., Okhovatian, S., Savoji, H., Campbell, S. B., Lai, B. F. L., Wu, J., Pascual-Gil, S., Bannerman, D., Rafatian, N., Li, R. K., & Radisic, M. (2022). Toward Hierarchical Assembly of Aligned Cell Sheets into a Conical Cardiac Ventricle Using Microfabricated Elastomers. Advanced Biology. 10.1002/adbi.202101165

Ng, E. S., Davis, R., Stanley, E. G., & Elefanty, A. G. (2008). A protocol describing the use of a recombinant protein-based, animal product-free medium (APEL) for human embryonic stem cell differentiation as spin embryoid bodies. Nature Protocols, 3(5), 768–776. 10.1038/nprot.2008.42

Nunes, S. S., Miklas, J. W., Liu, J., Aschar-Sobbi, R., Xiao, Y., Zhang, B., Jiang, J., Massé, S., Gagliardi, M., Hsieh, A., Thavandiran, N., Laflamme, M. A., Nanthakumar, K., Gross, G. J., Backx, P. H., Keller, G., & Radisic, M. (2013). Biowire: A platform for maturation of human pluripotent stem cell-derived cardiomyocytes. Nature Methods, 10(8), 781–787. 10.1038/nmeth.2524

Patel, N. M., & Birla, R. K. (2018). The bioengineered cardiac left ventricle. ASAIO Journal, 64(1), 56–62. 10.1097/MAT.0000000000000642

Pound, P., & Ritskes-Hoitinga, M. (2018). Is it possible to overcome issues of external validity in preclinical animal research? Why most animal models are bound to fail. In Journal of Translational Medicine (Vol. 16, Issue 1). BioMed Central Ltd. 10.1186/s12967-018-1678-1

Sacchetto, C., Vitiello, L., de Windt, L. J., Rampazzo, A., & Calore, M. (2020). Modeling cardiovascular diseases with hipsc-derived cardiomyocytes in 2d and 3d cultures. In International Journal of Molecular Sciences (Vol. 21, Issue 9). MDPI AG. 10.3390/ijms21093404

Schwach, V., Slaats, R. H., Cofiño-Fabres, C., ten Den, S. A., Rivera-Arbeláez, J. M., Dannenberg, M., van Boheemen, C., Ribeiro, M. C., van der Zanden, S. Y., Nollet, E. E., van der Velden, J., Neefjes, J., Cao, L., & Passier, R. (2024). A safety screening platform for individualized cardiotoxicity assessment. IScience, 27(3). 10.1016/j.isci.2024.109139

Takahashi, K., Tanabe, K., Ohnuki, M., Narita, M., Ichisaka, T., Tomoda, K., & Yamanaka, S. (2007). Induction of Pluripotent Stem Cells from Adult Human Fibroblasts by Defined Factors. Cell, 131(5), 861–872. 10.1016/j.cell.2007.11.019

Tsao, C. W., Aday, A. W., Almarzooq, Z. I., Anderson, C. A. M., Arora, P., Avery, C. L., Baker-Smith, C. M., Beaton, A. Z., Boehme, A. K., Buxton, A. E., Commodore-Mensah, Y., Elkind, M. S. V., Evenson, K. R., Eze-Nliam, C., Fugar, S., Generoso, G., Heard, D. G., Hiremath, S., Ho, J. E., … Martin, S. S. (2023). Heart Disease and Stroke Statistics - 2023 Update: A Report from the American Heart Association. In Circulation (Vol. 147, Issue 8, pp. E93–E621). Lippincott Williams and Wilkins. 10.1161/CIR.0000000000001123

Wei, A. E., Maslov, M. Y., Pezone, M. J., Edelman, E. R., & Lovich, M. A. (2014). Use of pressure-volume conductance catheters in real-time cardiovascular experimentation. Heart Lung and Circulation, 23(11), 1059–1069. 10.1016/j.hlc.2014.04.130

Yang, X., Pabon, L., & Murry, C. E. (2014). Engineering adolescence: Maturation of human pluripotent stem cell-derived cardiomyocytes. In Circulation Research (Vol. 114, Issue 3, pp. 511–523). 10.1161/CIRCRESAHA.114.300558

