## Supplementary Information for "A human engineered mini-heart platform for mimicking ventricular pump function"

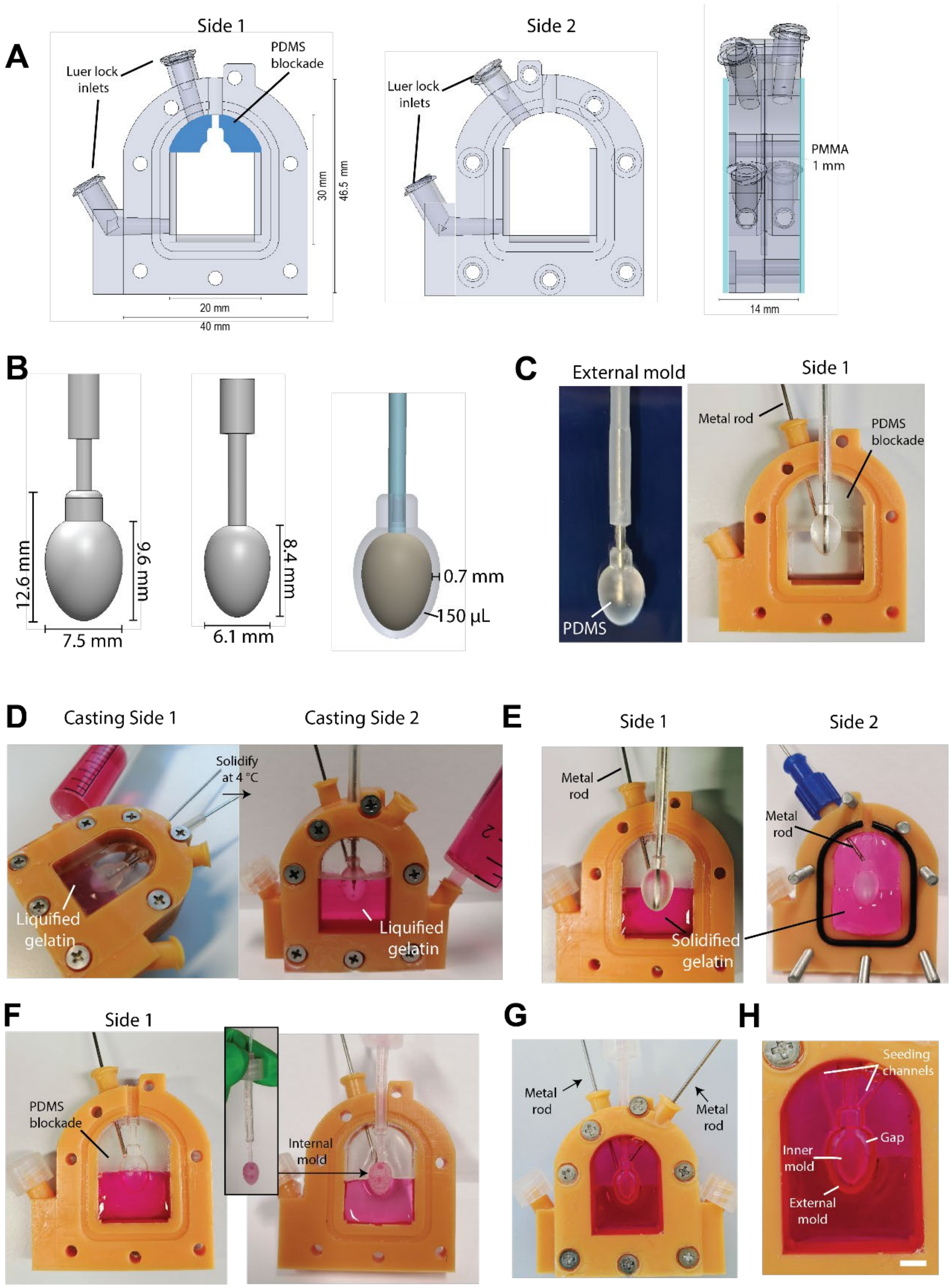


**Figure S1. Fabrication of mini-heart molds.** A) Design of custom 3D-printed bioreactor composed of 2 parts (Side 1 and Side 2). B) Dimensions of external (left) and internal (middle) molds, with their respective volume difference upon alignment (right). C) Insertion of PDMS external mold into the PDMS blockade on Side 1 of the bioreactor. D) Casting of gelatin around external PDMS mold inside the closed bioreactor. The process is done in 2 steps: first filling Side 1 horizontally, waiting for gelation at 4 °C, followed by filling of Side 2 with new liquified gelatin (right) until the bioreactor is completely filled. E) Removal of PDMS external mold from the open bioreactor once gelatin parts have solidified. F) Alignment of internal gelatin mold within the gelatin bodies casted inside the bioreactor G) Reassembly of bioreactor, establishing an ellipsoid gap between the gelatin bodies. Metal rods at luer lock inlets (arrows) are removed to create channels for seeding. H) Seeding channels and gap established between gelatin bodies (scale bar= 1 mm).

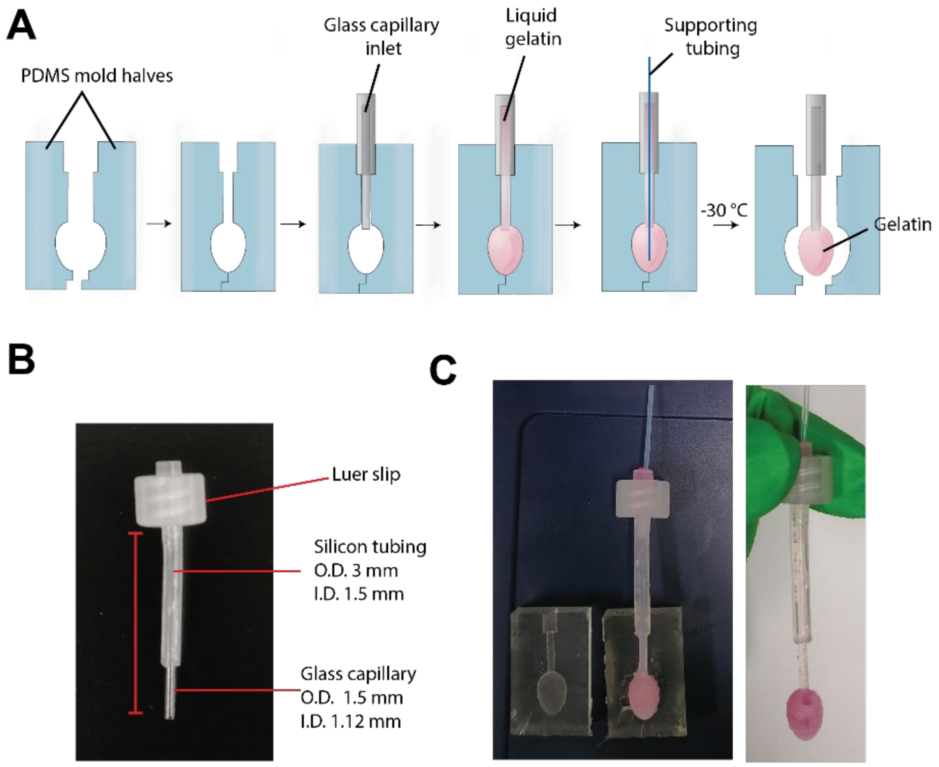


**Figure S2. Fabrication of mini-heart internal molds.** A) Schematic illustrating the casting process of the internal gelatin mold, guided by two PDMS halves that serve as negative mold. Liquified gelatin is introduced through the glass capillary inlet and subsequently frozen before removal from the PDMS halves. B) Dimensions of the glass capillary inlet, which is surrounded by a silicone tubing and connected to a luer slip. C) Removal of internal gelatin mold from PDMS halves.

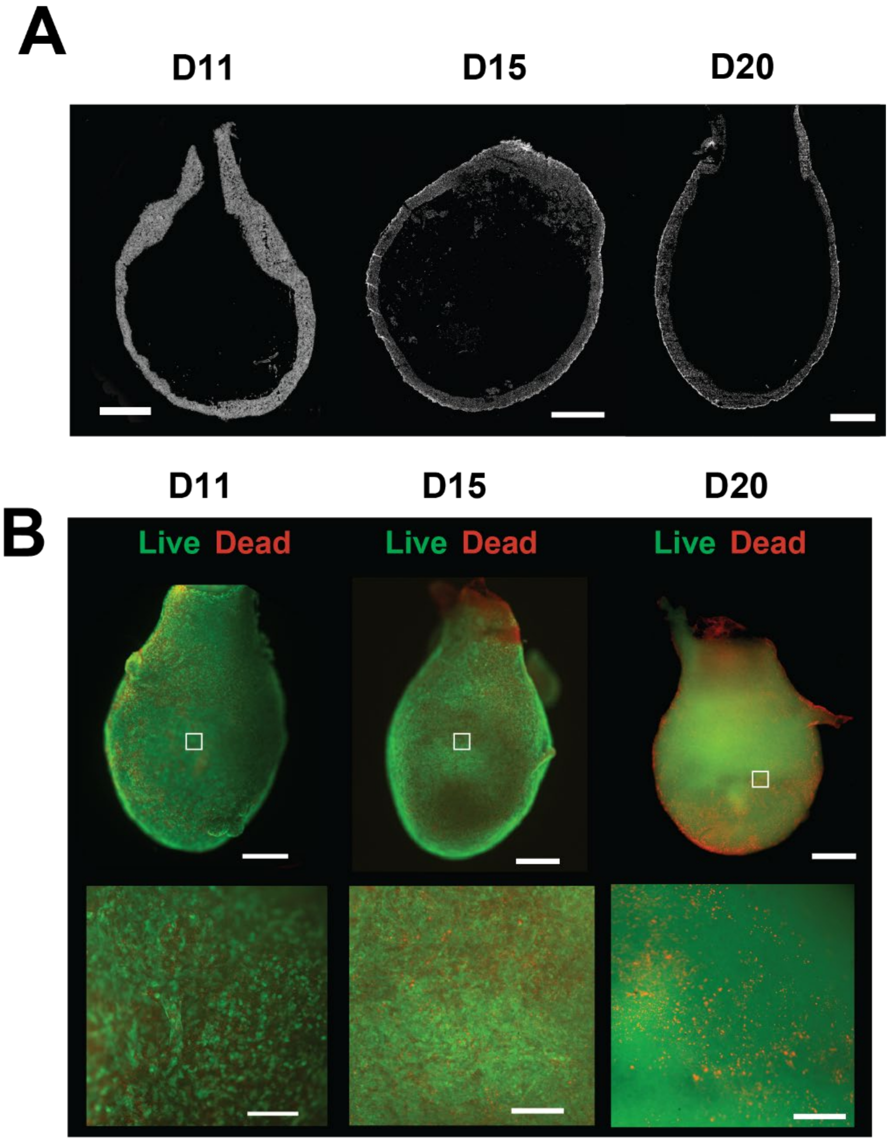


**Figure S3. Characterization of mini-hearts.** A) Frontal cryosections of mini-hearts at days 11, 15, and 20 (Scale bars= 1mm). B) Live-dead staining of mini-hearts at days 11, 15, and 20 (upper row scale bars= 1 mm; lower row scale bars= 300 μm).

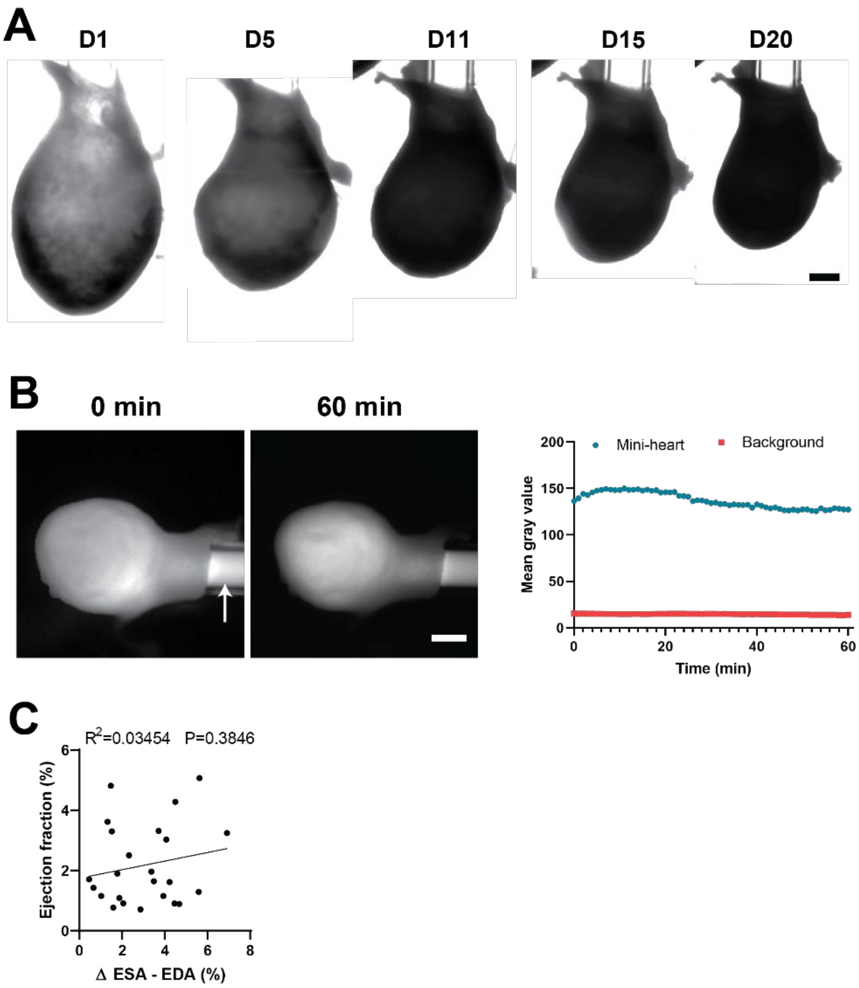


**Figure S4. Characterization of tissue compaction and permeability.** A) Progression of mini-heart compaction on days 1, 5, 11, 15, and 20 (scale bar= 1 mm). B) Injection of fluorescently-labelled dextran (40kDa) through mini-heart inlet (white arrow) at 0 and 60 minutes post-injection (Scale bar= 1 mm). The right panel displays corresponding mean gray values inside and outside the mini-heart over time. C) Correlation between relative end-systolic area change and ejection fraction.

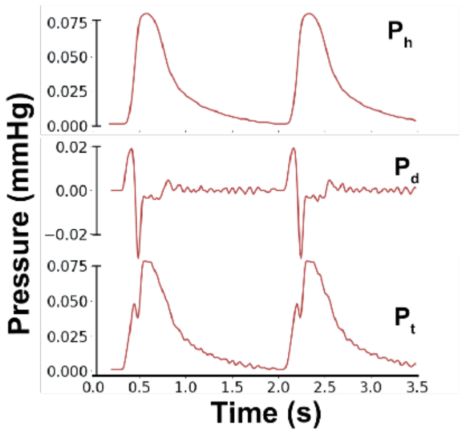


**Figure S5. Pressure developed during mini-heart beating cycles.** Estimated total pressure ($P_t$) inside the mini-heart in time, composed by the sum of hydrostatic ($P_h$) and dynamic ($P_d$) components.

**Supplementary video 1.** Magnified view of spontaneously contracting mini-heart at culture day 15. Scale bar: 1 mm.

**Supplementary video 2.** View of bioreactor and spontaneously beating of mini-heart. Mini-heart contraction is synchronized with liquid displacement in the glass capillary inlet. Scale bar: 1 cm.

**Supplementary video 3.** Magnified view of liquid displacement in glass capillary coupled to the inlet of the mini-heart during spontaneous beating. Scale bar: 1 mm.

**Supplementary video 4.** Magnified view of mini-heart beating spontaneously at Day 11. Scale bar: 3 mm.

**Supplementary video 5.** Magnified view of mini-heart beating spontaneously at Day 15. Scale bar: 3 mm.

**Supplementary video 6.** Magnified view of mini-heart beating spontaneously at Day 20. Scale bar: 3 mm.

**Supplementary video 7.** Calcium transient propagation of mini-heart at culture day 14. Scale bar: 1 mm.

.
